# Acute Degradation of Pumilio Proteins Uncovers a Biphasic Post-transcriptional Regulatory Hierarchy Controlling Embryonic Stem Cell Fate Decisions

**DOI:** 10.64898/2026.03.10.710821

**Authors:** Yuedong Huang, Wenxue Li, Hanna Richman, Yansheng Liu, Haifan Lin

## Abstract

Post-transcriptional regulation is critical for mammalian embryogenesis yet has been underexplored. We previously showed that RNA-binding Pumilio proteins (Pum1/2) are essential for early mouse embryogenesis and embryonic stem cell (ESC) functions. Here, using acute protein degradation systems combined with time-resolved RNA-seq and eCLIP, we delineate a two-phase regulatory hierarchy modulated by Pum1/2 in mouse ESCs. The first phase, occurring within 10 hours of Pum1/2 depletion, is predominantly the stabilization of over 100 Pum1/2-target mRNAs, while the second phase, occurring in subsequent 66 hours, propagates to over 1,000 mRNAs mostly through indirect regulatory effects. Functionally, Pum1/2 depletion delays transition from naïve to formative pluripotency, impairs neuroectoderm differentiation, and enhances germline specification. Mechanistically, Pum1/2 directly repress mRNAs encoding PRC2 subunits, including *Suz12*, thereby constraining H3K27me3 deposition at neuroectodermal gene loci. These findings establish Pum1/2 as biphasic post-transcriptional regulators of pluripotency and lineage balance and link RNA stability control to chromatin-mediated silencing.

**HIGHLIGHTS:**

- Acute Pum1/2 degradation provides temporal resolution for profiling post-transcriptional regulation.
- Pum1/2 destabilize more than 100 direct targets and modulate a biphasic network of over 1,000 mRNAs.
- Pum1/2 loss delays pluripotency transition, suppresses neuroectoderm, and promotes germline fate.
- Pum1/2 directly regulate *Suz12* mRNA decay to modulate PRC2-mediated repression.

## INTRODUCTION

Mammalian embryogenesis is a highly orchestrated process that requires regulation across epigenetic^1,2^, transcriptional^3,4^, and post-transcriptional levels^5,6^. As embryos progress from blastocyst to gastrulation, these regulatory networks are extensively rewired to ensure precise control of cell fate divergence and embryonic patterning^1^. During this process, the naïve pluripotency circuit in epiblasts is disassembled, followed by activation of lineage specification signaling and generation of three distinct germ layers and primordial germ cells (PGCs)^7–11^. While the roles of epigenetic and transcriptional regulators in this transition have been extensively characterized^8,10,12,13^, the contribution of post-transcriptional regulation is much less studied.

Pum1 and Pum2, the two murine Pumilio-family RNA-binding proteins, are essential for early embryogenesis and embryonic stem cell (ESC) functions^14–16^. *Pum1* knockdown in haploid mouse ESCs (mESCs) delayed pluripotency exit, due to stabilization of naïve pluripotency factors directly bound and regulated by Pum1, including *Sox2*, *Tbx3*, *Esrrb*, *Tfcp2l1*, and *Klf2*^14^. *Pum1/2* double-knockout (DKO) in mice caused delayed morula-to-blastocyst transition, compromised blastocyst development, and lethality by embryonic day (E) 8.5^15–17^. The mechanisms underlying this lethality remain unclear, but it has been proposed that loss of Pum1 and Pum2 led to premature primitive endoderm (PrE) differentiation driven by de-repression of *Gata6*, a key PrE regulator and a direct Pum target^15^. Uyhazi and Yang et al. further reported that Pum1 and Pum2 have distinct functions in mESCs, with Pum1 promoting differentiation and Pum2 supporting self-renewal, respectively^16^. Because these studies^14–16^ used RNAi or gene knockout to deplete Pum1/2 days or even earlier before transcriptome analysis, and mESCs could have adapted to this perturbation over time, they could not establish a temporal hierarchy of regulatory events among targets, thereby limiting mechanistic insight into Pum1/2 function.

Recent advances in targeted protein degradation technologies, such as the HaloPROTAC and dTAG systems, have enabled rapid and inducible protein depletion in cells within hours, providing a powerful tool for studying gene regulatory effects with high temporal resolution^18,19^. Leveraging these technologies, we generated mESCs in which endogenous *Pum1* and *Pum2* genes were tagged at the start of their open reading frames (ORFs) with HaloTag- and FKBP12^F36V^-coding sequences, respectively. Using these mESCs, we performed temporally controlled depletion of Pum1 and/or Pum2 with small-molecule PROTAC degraders, followed by time-course RNA-seq during mESC differentiation. This approach, combined with enhanced crosslinking and immunoprecipitation (eCLIP) analysis of Pum1/2-binding sites in mESCs, enabled unambiguous identification of over 100 mRNAs that are directly bound and regulated by Pum1 and Pum2 at the level of RNA stability. In dual Pum1/2-depleted mESCs, stabilization of these direct regulatory targets led to deregulation of over 1,000 mRNAs during differentiation, resulting in gene expression changes indicative of delayed naïve-to-formative transition, reduced neuroectoderm (NE) differentiation, and enhanced PGC specification. Pum1/2 achieve these functions in part by down-regulating mRNAs encoding PRC2 subunits, including *Suz12*. Loss of Pum1/2 increases PRC2 activity and H3K27me3 levels during mESC differentiation, thereby repressing NE regulators. Together, our findings establish Pum1 and Pum2 as key post-transcriptional regulators of early cell fate decisions and reveal a novel link between Pum1/2-mediated mRNA decay and PRC2-mediated epigenetic silencing.

## RESULTS

### HaloPROTAC3 and dTAG-13 enable rapid and orthogonal depletion of Pum1 and Pum2 in *Pum1*Halo/Halo; *Pum2*dTAG/dTAG mESCs

To investigate the direct regulatory roles of Pum proteins in mESCs, we employed two orthogonal targeted protein degradation systems – HaloPROTAC and dTAG – to achieve rapid and specific depletion of Pum1 and Pum2, respectively (Figure 1A). Structural predictions using PrDOS^20^ and AlphaFold^21,22^ indicated that the N-termini of Pum1 and Pum2 are intrinsically disordered (Figure S1A), suggesting that modifications in these regions are unlikely to disrupt protein functions. Therefore, we used CRISPR-mediated knock-in to fuse a 3xFLAG-HaloTag-coding sequence to the start of *Pum1* ORF and 2xHA-FKBP12^F36V^-coding sequence to the start of *Pum2* ORF in mESCs (Figure 1B). Genotyping of clonal lines confirmed successful editing of both alleles of *Pum1* and *Pum2* (Figures S1B and S1C). The resulting homozygous double knock-in line were designated as *Pum1*^Halo/Halo^; *Pum2*^dTAG/dTAG^ mESCs. Immunoblot analysis of a representative clone, 2F4, verified expression of the tagged Pum proteins, with Halo-Pum1 expressed at ∼2x the level of endogenous Pum1 and FKBP12^F36V^-Pum2 expressed at levels comparable to endogenous Pum2 (Figures 1C and 1D), indicating that both fusion proteins are stable in mESCs in the absence of their respective degraders. Based on these results and unaltered pluripotency of 2F4 mESCs (see below), this clone and its derivatives were used for subsequent experiments.

**Figure 1.**
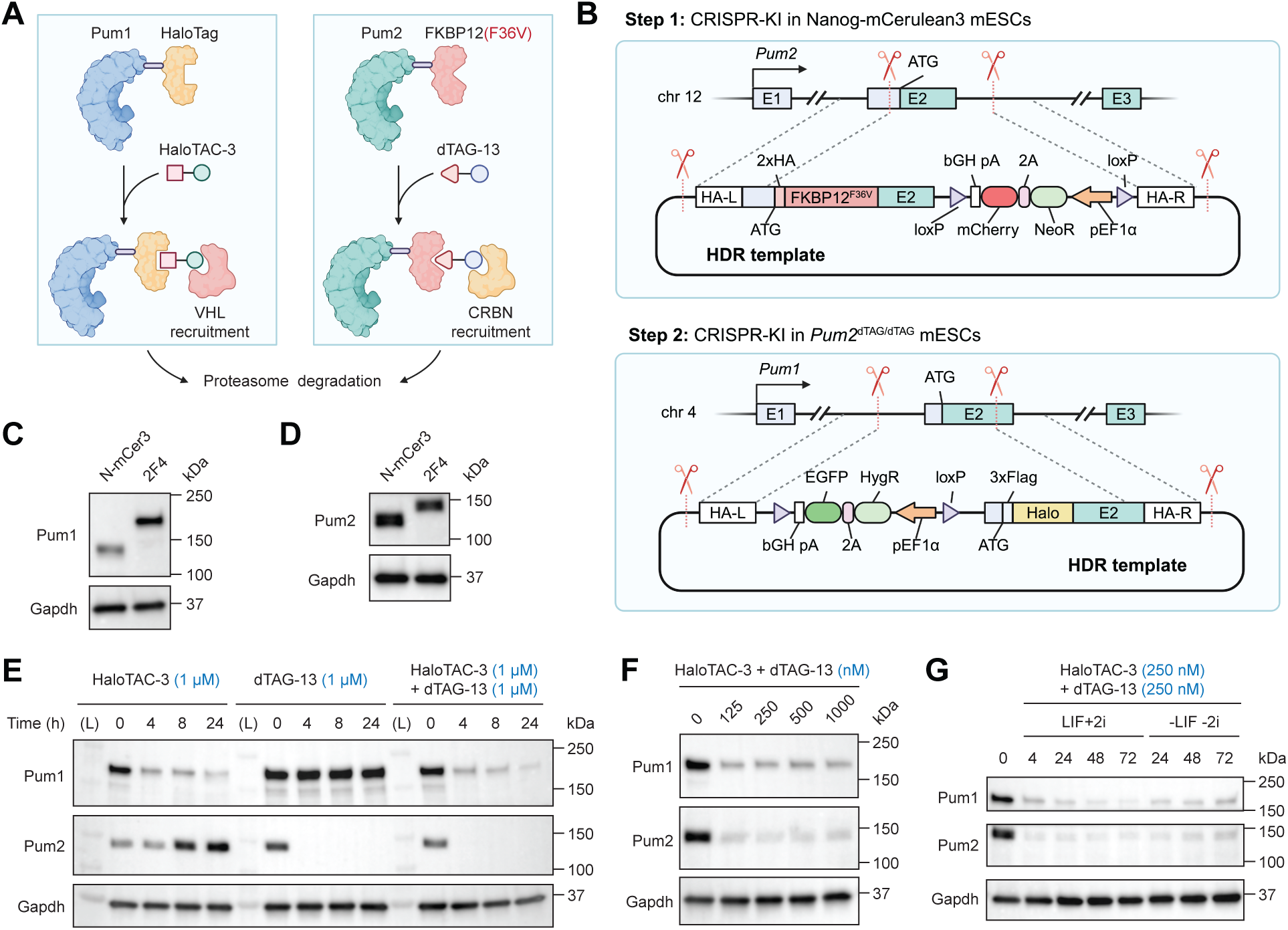
PROTAC-based inducible degradation of Pum1/2 in mESCs. (A and B) Schematic illustrations of (A) the HaloPROTAC and dTAG systems for orthogonal degradation of Halo-Pum1 and/or FKBP12^F36V^-Pum2, and (B) the CRISPR knock-in strategy. Schematics were created with BioRender. (C and D) Western blots of mESCs before (N-mCer3) and after (2F4) CRISPR-KI, comparing expression levels of (C) Halo-Pum1 versus wild-type Pum1 and (D) FKBP12^F36V^-Pum2 versus wild-type Pum2. (E–G) Western blots of 2F4 mESCs treated with the indicated concentrations of PROTACs for the indicated time course (F: LIF+2i, 4 h).

To evaluate the efficiency and specificity of the degradation systems, we treated 2F4 mESCs with 1 µM HaloPROTAC3 (also referred to as HaloTAC-3), 1 µM dTAG-13, or both compounds for 24 hours (Figure 1E). HaloTAC-3 specifically and significantly reduced Halo-Pum1 levels within four hours of treatment and, notably, led to upregulation of FKBP12^F36V^-Pum2 levels by eight hours. This increase was sustained at 24 hours post-treatment. This observation is consistent with previous reports that Pum1 negatively regulates *Pum2* mRNA stability^16,23^. Furthermore, it reveals that Pum1 requires more than four but fewer than eight hours for its initial regulatory effect on target mRNAs to manifest at the protein level. This finding supports our choice of four hours as the earliest post-treatment time point for RNA-seq analysis (see below), as it is sufficiently early to capture the onset of Pum1-mediated regulation. Similarly, dTAG-13 efficiently depleted FKBP12^F36V^-Pum2 within four hours without affecting Halo-Pum1 levels, indicating both the specificity of dTAG-induced degradation and the minimal impact of Pum2 loss on Pum1 expression (Figure 1E). Finally, combined treatment with HaloTAC-3 and dTAG-13 resulted in near-complete degradation of both Pum1 and Pum2, confirming the effectiveness of the PROTAC degradation systems (Figure 1E).

To minimize potential off-target effects, we titrated both compounds and determined that 125 nM of either HaloTAC-3 or dTAG-13 was sufficient for maximal target degradation within 4 hours (Figure 1F). We selected 250 nM as the working concentration for all subsequent experiments, as this dosage achieved sustained depletion of Pum proteins in mESCs for at least 72 hours in both naïve pluripotency and differentiation conditions without medium change (Figure 1G).

Together, these results establish a robust and inducible system for rapid, orthogonal protein degradation, expanding the current repertoire of combinatorial degron tools^24^. This platform provides precise temporal control to dissect the individual and combined regulatory roles of Pum1 and Pum2 during pluripotency transitions and cell fate decisions of mESCs.

### Pum1 and Pum2 regulate the stability of over 100 target mRNAs within first 10 hours

Previous studies using siRNA-based knock-down or Cre-loxP-mediated knockout were not able to temporally resolve the sequence of regulatory events following Pum1/2 depletion in ESCs, mostly due to the early depletion of Pum proteins that occurred days or even earlier before the examination time^14,16^. To fill in this gap, we designed a time-course RNA-seq based on the fast-acting HaloPROTAC and dTAG systems. First, we performed Pum1 and/or Pum2 depletion in mESCs for 4 hours and then, with continuous HaloTAC-3 and/or dTAG-13 presence, either maintained the cells under naïve pluripotency condition or started spontaneous differentiation by LIF and 2i withdrawal (Figure 2A). Cells were collected at zero and six hours following the four-hour Pum1/2 depletion for RNA-seq. Differential gene expression analysis was performed using an adjusted p-value cutoff of 0.01 and a log_2_ fold-change (log_2_FC) threshold of ±0.3 to match the regulatory effect of a single Pumilio Response Element (PRE), as reported by Bohn et al.^25^ This threshold corresponds to approximately a 1.25-fold increase or 0.8-fold decrease in mRNA levels for up- and down-regulated genes, respectively.

**Figure 2.**
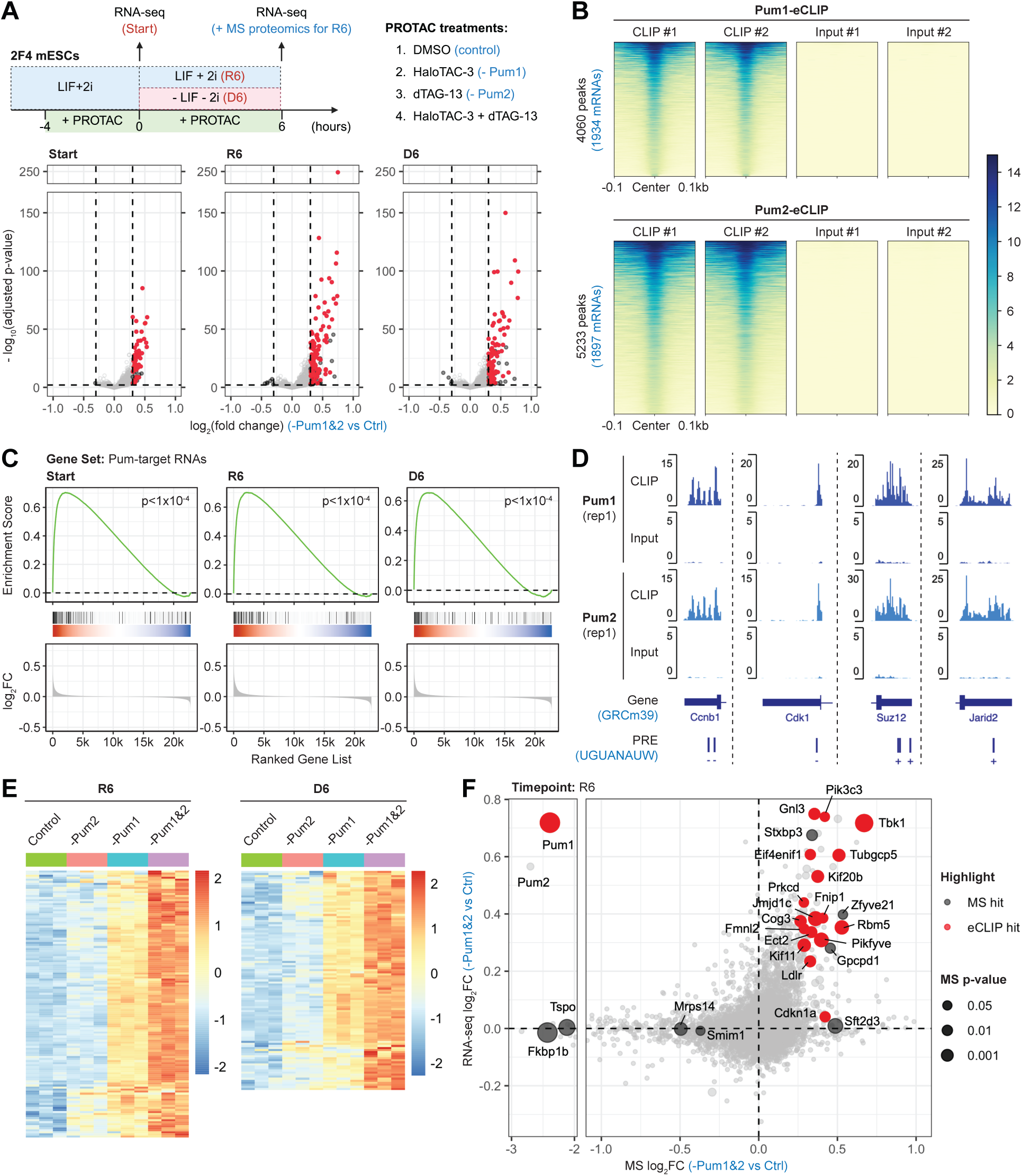
Pum1/2 depletion leads to immediate upregulation of a subset of their direct target mRNAs. (A) Design of the time-course RNA-seq experiment (upper panel) and volcano plots of differentially regulated RNAs at the indicated time points (lower panels). Significance cutoffs: |log_2_FC| > 0.3, adjusted p-value < 0.01. The direct Pum1/2 targets are highlighted in red. (B) Heatmap of significant Pum1/2-eCLIP peaks, with corresponding read coverage of input samples shown in parallel. (C) GSEA of Pum1/2-target mRNAs in the DEGs upon dual Pum1/2-depletion versus control. (D) Pum1/2-eCLIP read-coverage tracks for the selected target mRNAs. The genomic strand on which the PRE elements reside is indicated by “+” or “–”. (E) Heatmaps showing DESeq2-normalized counts of the DEGs in 2F4 mESCs with indicated treatments at R6 and D6. Colors represent the z-scores (normalized to row means). (F) Scatter plot showing log_2_FC of protein abundance in DIA-MS and the corresponding log_2_FC of RNA abundance in RNA-seq (dual Pum1/2-depletion vs. control, R6). Significantly changed proteins in DIA-MS (adjusted p-value < 0.05) are highlighted in black, and the Pum1/2-target RNAs among them are further highlighted in red. Circle sizes inversely represent the p-values in DIA-MS.

RNA-seq analysis revealed a striking bias toward upregulation of mRNAs following Pum1/2 depletion in mESCs (Figure 2A). At four hours of depletion (Start), we identified 61 significantly upregulated mRNAs and only one mildly downregulated. Similarly, in the next six hours of culture under either the self-renewal (R6) or differentiation (D6) condition, 131 and 107 mRNAs were upregulated, respectively, while only six and four mRNAs were downregulated (Table S1).

To determine whether Pum proteins directly bind to the upregulated mRNAs, we performed eCLIP in naïve mESCs and identified 1,934 direct mRNA targets of Pum1 and 1,897 of Pum2 (peaks significantly enriched over input: log_2_FC > 4.5, p < 1x10^-10^), with 1,368 targets shared between the two proteins (Figure 2B and Table S2). To further assess the similarity of their binding profiles, we quantified read coverage of Pum1- and Pum2-eCLIP across both Pum1 and Pum2 binding clusters and observed very strong correlations (Pearson *r* = 0.95-0.97), indicating that Pum1 and Pum2 bind to nearly identical sites across a largely overlapping set of target mRNAs (Figures S2A and S2B). As expected, *de novo* motif search using the HOMER algorithm identified the canonical PRE (UGUANAUA) as the top and only reproducible eight-nucleotide binding motif for both proteins (Figure S2C). Given the high similarity between their RNA-binding profiles, we combined the 1,934 Pum1-target and 1,897 Pum2-target mRNAs, resulting in a total of 2,463 Pum1/2-target mRNAs in mESCs (Table S2).

By integrating eCLIP and RNA-seq data, we found that the upregulated mRNAs following Pum1/2 depletion were significantly enriched for direct Pum1/2 targets identified by eCLIP (Figure 2A). Specifically, 49 out of 61 (80%) upregulated mRNAs at 4 hours post-depletion were direct Pum1/2 targets. At R6 and D6, 99 out of 131 (76%) and 79 out of 107 (74%) upregulated mRNAs, respectively, were direct Pum1/2 targets (Figure 2A). Comparison of these upregulated transcripts revealed substantial overlap across time points, indicating a core set of mRNAs consistently destabilized by Pum proteins (Figure S2D). Gene set enrichment analysis (GSEA) considering differential regulation of all genes regardless of fold-change thresholds further confirmed the significant enrichment of direct Pum1/2 targets among the upregulated mRNAs following dual Pum1/2 depletion (Figure 2C).

Overall, the immediate upregulation of direct Pum1/2-target mRNAs following acute Pum1/2 depletion is consistent with the well-established role of Pum proteins in promoting target RNA degradation^26^. In contrast, only a small number of mRNAs were downregulated at R6 and D6, and none of them were eCLIP targets of Pum1/2 (Figure 2A), indicating that these effects are likely indirect. Thus, transcriptomic changes observed within 10 hours of PROTAC treatment largely reflect direct regulatory effects of Pum proteins, with the majority of upregulated mRNAs being their direct targets. These findings underscore the advantage of acute protein degradation by our system over RNAi or gene knockout approaches, as it provides greater temporal resolution for distinguishing direct from indirect regulatory consequences.

Gene Ontology (GO) analysis of differentially expressed genes (DEGs) at R6 and D6 (log_2_FC > 0, adjusted p < 0.01) revealed enrichment in diverse biological processes, including regulation of cell cycle process, transcription by RNA Polymerase II, and protein phosphorylation, indicating that Pum proteins regulate a broad range of cellular functions (Figure S2E). These direct Pum1/2-target mRNAs include cell cycle regulators such as *Ccnb1* and *Cdk1*, as well as epigenetic factors that regulate ESC fate decisions, including PRC2 components *Suz12* and *Jarid2* (Figure 2D).

Compared to double Pum1/2 depletion, degradation of Pum1 or Pum2 alone resulted in significantly fewer upregulated mRNAs (Figure S2F), likely due to functional compensation between the two proteins^16,23,25^. Notably, Pum1 depletion resulted in a higher magnitude of mRNA upregulation than Pum2 depletion, likely due to the higher abundance of Pum1 protein in mESCs, as indicated by intensity-based absolute quantification (iBAQ) values from data-independent acquisition mass spectrometry (DIA-MS) proteomics (Table S3). Despite this difference, transcriptomic changes following Pum1 or Pum2 depletion were moderately correlated under both self-renewal and differentiation conditions (Figure S2F). Furthermore, additional depletion of Pum2 in Pum1-depleted mESCs enhanced the upregulation of DEGs (Figure 2E). These findings indicate that (1) Pum1 and Pum2 largely regulate the same set of RNAs, with Pum1 exerting stronger regulatory effects; and (2) both of them act as negative regulators of RNA stability, as reflected by the predominant upregulation of mRNAs following Pum1 and/or Pum2 depletion.

To assess whether Pum proteins also regulate translation of their target mRNAs, we performed total cellular proteomic profiling of R6 samples using DIA-MS. Proteomic analysis confirmed efficient depletion of Pum1 and/or Pum2 proteins by PROTAC treatment (Figure 2F). Following double Pum1/2 depletion, the majority of upregulated proteins corresponded to mRNAs that also showed increased abundance with similar levels of fold-changes, indicating that these changes were largely, if not exclusively, due to enhanced mRNA stability rather than increased translation (Figure 2F). One notable exception was *Cdkn1a*, a well-characterized Pum1/2 target, which showed a significant increase in protein level with minimal change in mRNA abundance following Pum1/2 depletion (Figure 2F), indicating that it is primarily regulated at the translational level. These results indicate that in mESCs, Pum proteins predominantly act by destabilizing their target RNAs, with a smaller subset of targets – such as *Cdkn1a* – being regulated mainly at the level of translation.

### Pum1 and Pum2 regulate a second phase of gene expression of over 1,000 mRNAs involved in ESC pluripotency and fate decisions within subsequent 66 hours

Having characterized the direct regulatory effects of Pum1 and Pum2 on their target mRNAs across diverse cellular pathways, we next examined the transcriptomic responses to Pum1/2 depletion over an extended time course. To this end, 2F4 mESCs were treated with PROTACs for 4 hours under LIF/2i condition to deplete Pum1 and/or Pum2, followed by differentiation for 6, 24, or 72 hours with continuous PROTAC treatment (Figure 3A). A parallel RNA-seq was performed on mESCs maintained under LIF/2i to distinguish differentiation-specific responses to Pum1/2 depletion.

**Figure 3.**
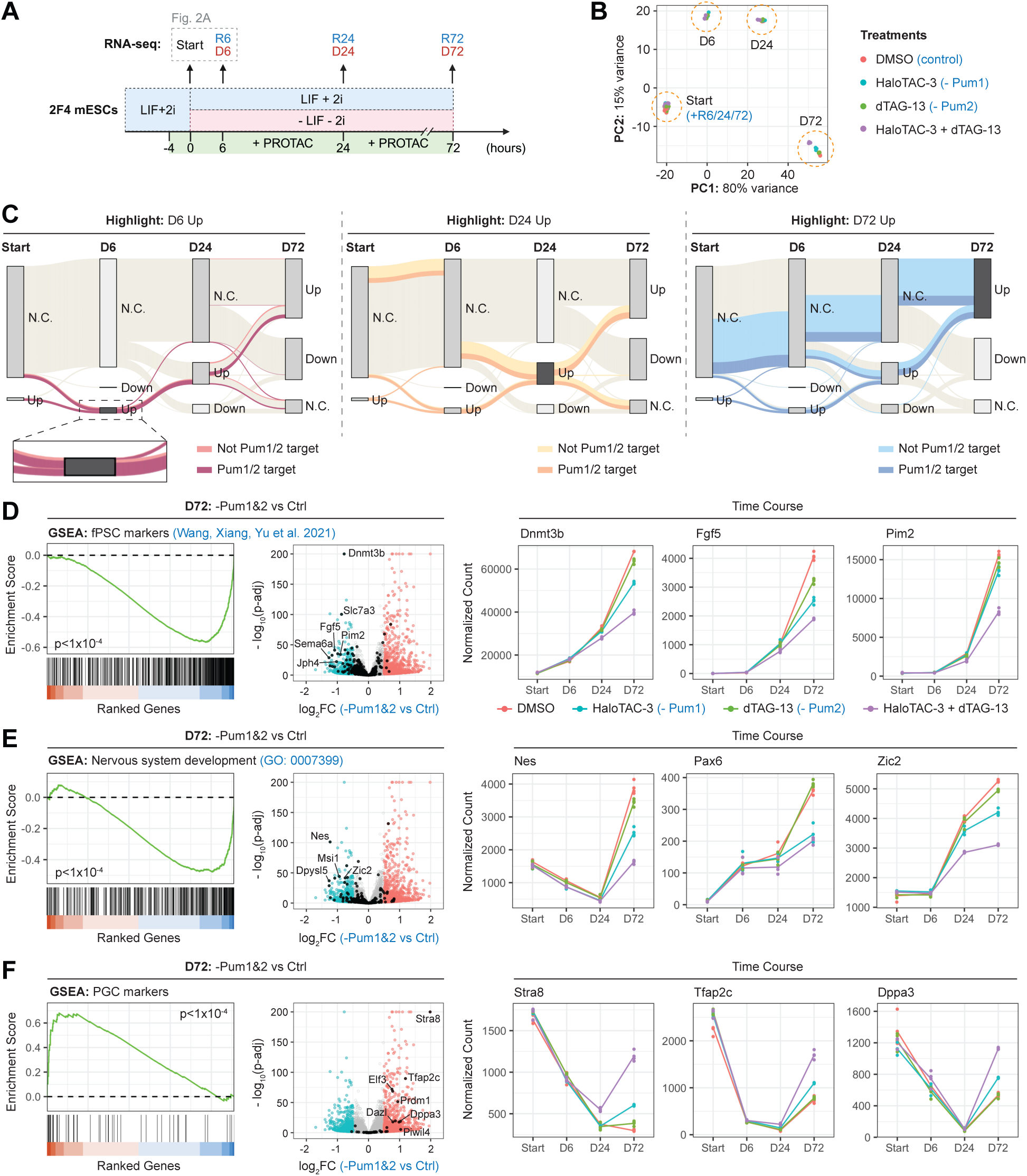
Pum1/2 regulate pluripotency transitions and cell fate decisions of mESCs. (A) Design of the time-course RNA-seq experiment. Four PROTAC treatment groups: (1) DMSO (control), (2) HaloTAC-3, (3) dTAG-13, and (4) HaloTAC-3 + dTAG-13. (B) Principal component analysis (PCA) of RNA-seq data. (C) Sankey diagrams showing the trajectories of upregulated DEGs (log_2_FC > 0.3, adjusted p-value < 0.01) at D6, D24, and D72. Pum1/2 targets and non-targets are distinguished by line colors. N.C. indicates genes with no change in expression. (D–F) GSEA and corresponding volcano plots for (D) fPSC markers, (E) nervous system development-related genes, and (F) PGC markers at D72, comparing Pum1/2 double-depletion versus control. The specific gene sets are highlighted in black. Trend plots show mRNA levels (DESeq2-normalized counts) of the indicated genes during mESC differentiation (n = 3).

To globally evaluate the impact of Pum1/2 depletion on gene expression dynamics, we performed principal component analysis (PCA) on the RNA-seq data (Figure 3B). The first principal component (PC1) separated samples by culture condition (self-renewal vs. differentiation), reflecting the extensive transcriptomic changes during ESC differentiation. At D72, distinct clustering of different PROTAC treatment groups indicated that depletion of Pum1 and/or Pum2 indeed significantly altered differentiation trajectories of ESCs in a similar fashion (Figure 3B).

When further assessing individual contributions of Pum1 and Pum2 loss, we found that transcriptomic changes following double depletion greatly exceeded the combined effects of single-Pum depletion (Figure S3A), consistent with the notion that Pum1 and Pum2 have overlapping functions and are able to partially compensate for each other when one is disrupted^16,23,25^. In support of this notion, additional depletion of Pum1 or Pum2 in cells lacking the other resulted in highly overlapping sets of DEGs at D72 (Figure S3B). Furthermore, Pum1/2 double depletion amplified gene expression changes observed in single-Pum depletion: mRNAs upregulated upon Pum1 or Pum2 loss were further upregulated in dual Pum1/2-depleted cells, and vice versa (Figure S3C). Together, these results indicate that Pum1 and Pum2 exert largely overlapping regulatory functions and are therefore likely to fulfill similar biological roles during ESC differentiation. This contrasts with the report by Uyhazi and Yang et al., who observed substantially different transcriptomes in Pum1- versus Pum2-KO mESCs and proposed distinct roles for Pum1 in promoting ESC differentiation and for Pum2 in sustaining self-renewal^16^. Future studies with extended time course up to 20 days (under differentiation or self-renewal conditions) will help determine whether functional divergence emerges at later stages following Pum1/2 loss.

We then focused on the Pum1/2-double-depleted mESCs, which showed the most pronounced transcriptomic changes during differentiation. We found that the proportion of direct Pum1/2 targets among upregulated DEGs steadily declined – from 80.0% at D6 to 26.2% at D24, and 13.9% at D72 – while substantial numbers of downregulated DEGs emerged from D24 onward (Figures 3C, S3A and S3D). This temporal pattern suggests a two-phase regulatory hierarchy modulated by Pum1/2: the first phase (Start and D6) marked by direct destabilization of >100 target mRNAs, followed by the second phase (D24 and D72) affecting >1,000 mRNAs largely through indirect mechanisms.

To identify differentiation programs shaped by this network, we performed GSEA on D72 samples using pluripotency gene sets (naïve, formative, primed) defined by Wang, Xiang, Yu et al.^27^ and cell-state markers from scRNA-seq studies of mouse gastrulation^8,13^. Notably, we did not observe significant delays in downregulation of naïve pluripotency factors within 72 hours of LIF and 2i withdrawal in Pum1/2-depleted mESCs as previously reported^14^ (Figure S3E). Instead, *Oct4* (*Pou5f1*) levels declined more rapidly between D6 and D72 (Figure S3E). These findings indicate that additional mechanisms beyond Pum1/2-mediated degradation of naïve pluripotency transcripts underlie the altered differentiation trajectory of Pum1/2-depleted mESCs.

Consistent with this, GSEA revealed significant downregulation of formative pluripotent stem cell (fPSC) markers in Pum1/2-double-depleted mESCs at D72, indicating a delayed transition from naïve to more advanced pluripotency states (Figure 3D). Examination of individual fPSC markers further confirmed their delayed induction during differentiation upon Pum1/2 loss (Figures 3D and S3E).

Since the formative pluripotency state serves as an important intermediate stage for subsequent germ layer specification^28^, delayed pluripotency transition may impair lineage commitment. Supporting this, we observed reduced differentiation toward the NE lineage in Pum1/2-double-depleted mESCs, as evidenced by significant downregulation of genes related to nervous system development in GSEA (Figure 3E). Expression dynamics of key NE markers confirmed an overall delay in ectoderm differentiation (Figures 3E and S3F). Additionally, GSEA revealed significant downregulation of nascent mesoderm and primitive streak markers (Figure S3G), indicating impaired differentiation toward these embryonic lineages following Pum1/2 depletion.

In contrast, marker genes for PGCs, including *Stra8*, *Tfap2c*, *Dppa3*, *Prdm1* (a.k.a. *Blimp1*), and *Elf3*, were upregulated in mESCs depleted of both Pum1 and Pum2 at D72, suggesting enhanced differentiation toward the PGC lineage (Figure 3F). Moreover, increased expression of parietal endoderm (Figure S3F) and extra-embryonic ectoderm markers (Figure S3G) reveals a potential bias in differentiation toward these extra-embryonic lineages after Pum1 and Pum2 depletion.

Overall, our findings define a two-phase regulatory hierarchy governed by Pum1/2 and highlight its essential roles in ensuring balanced germ layer differentiation of ESCs, a key feature of pluripotency. The biased lineage specification observed in Pum1/2-double-depleted mESCs may represent a major mechanism underlying the embryonic lethality of *Pum1/2*-DKO embryos *in vivo*.

### Pum1 and Pum2 regulate PRC2 mRNAs to balance cell fate decisions of ESCs

To identify direct Pum1/2 targets that act as intermediate regulators transmitting the effects of Pum1/2-depletion to indirect targets and thereby suppressing NE differentiation, we performed GSEA of transcription factor (TF) target genes^29,30^ in D72 and R72 samples, focusing on TFs whose mRNAs are direct Pum1/2 targets (Figures 4A and S4A). Notably, Suz12 target genes were significantly enriched among the downregulated transcripts in Pum1/2-double-depleted mESCs at D72 (Figure 4A). Suz12 is a core subunit of PRC2, which catalyzes H3K27 methylation to silence transcription of target genes. Our eCLIP and time-course RNA-seq confirmed that *Suz12* mRNA is a direct Pum1/2 target and is immediately upregulated following Pum1/2 depletion in mESCs (Figures 2D and 4B). This upregulation occurred under both self-renewal and differentiation conditions and persisted for at least 72 hours – the longest time point examined (Figure 4B). Importantly, GSEA enrichment of Suz12 target genes was specific to differentiation, as no significant enrichment was detected at R72 (Figure S4A). This indicates that Suz12 might play a more critical role during ESC differentiation than in naïve pluripotency maintenance, consistent with the previous studies showing that PRC2 is dispensable for maintaining ESC identity but essential for proper differentiation^31–33^.

**Figure 4.**
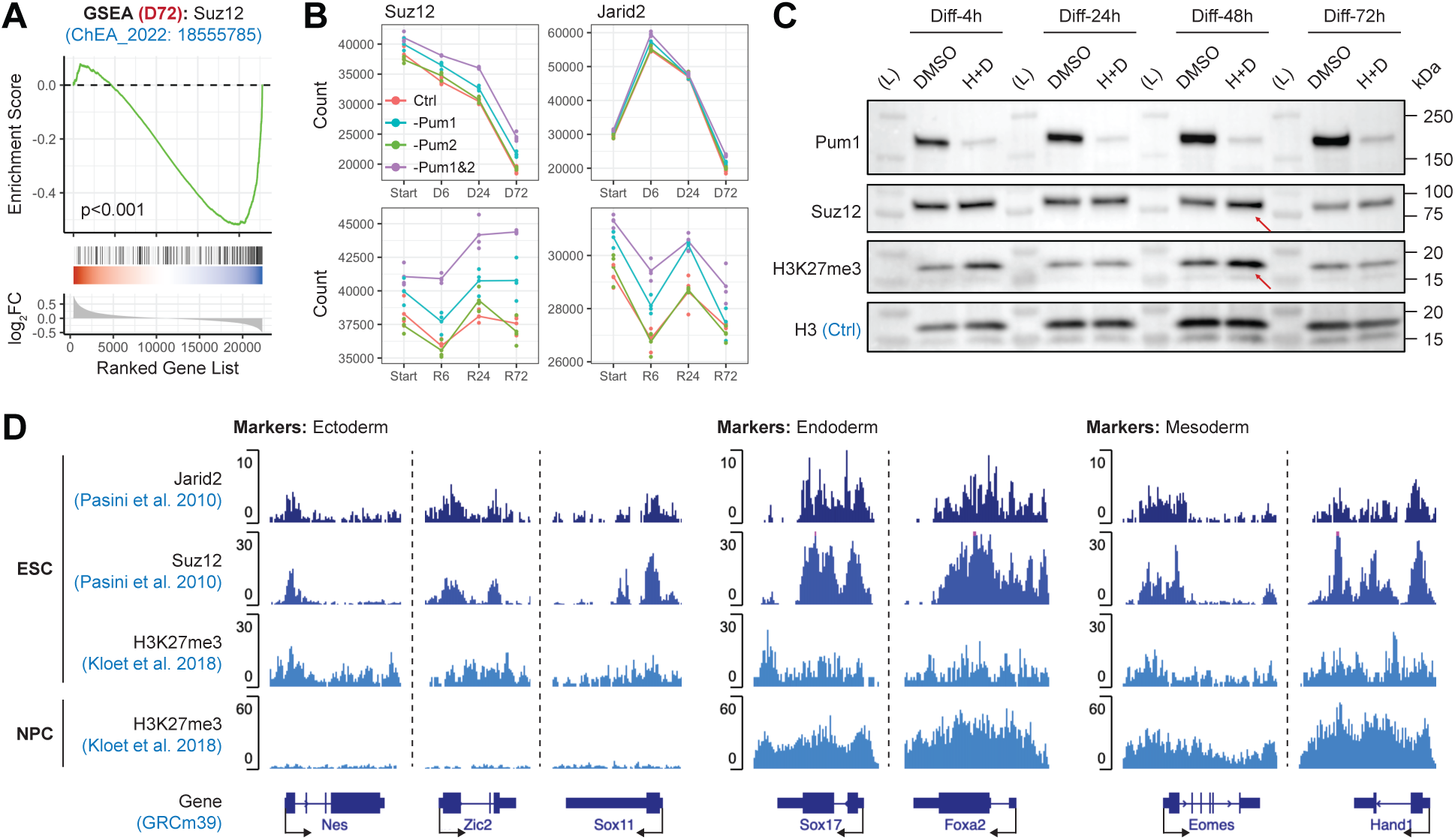
Pum1/2 regulate PRC2 mRNAs to balance cell fate decisions of mESCs. (A) GSEA for Suz12-target genes at D72 (dual Pum1/2-depletion vs. control). (B) Trend plots showing mRNA levels (DESeq2-normalized counts) of Suz12 and Jarid2 during mESC differentiation and self-renewal (n = 3). (C) Immunoblot analysis of Suz12 and H3K27me3 levels during differentiation of 2F4 mESCs treated with HaloTAC-3 and dTAG-13 (H+D) or with DMSO as controls. (D) ChIP-seq read density tracks for Suz12, Jarid2, H3K27me3 in ESCs and NPCs at ectoderm, endoderm and mesoderm markers.

Additionally, several other PRC2 subunits, including *Jarid2*, *Eed*, and *Rbbp7*, were also direct Pum1/2 targets and showed upregulation following Pum1/2 depletion, though to a lesser extent than *Suz12* (Figures 2D, 4B, and S4B). Meanwhile, *Kdm6b*, a histone demethylase that removes H3K27me3 marks, was downregulated in Pum1/2-depleted mESC during differentiation (Figure S4B). These results suggest that loss of Pum1/2 leads to upregulation of PRC2 components, potentially enhancing H3K27 methylation and repressing PRC2 target genes, thereby impairing NE differentiation.

To test this, we examined global H3K27me3 levels and found a significant increase within 48 hours of Pum1/2 depletion under differentiation conditions (Figure 4C). Analysis of published ChIP-seq datasets for Suz12^34^, Jarid2^34^ and H3K27me3^35^ in mESCs revealed that PRC2 and H3K27me3 marks are enriched at several NE marker genes, including *Nes*, *Zic2*, and *Sox11* (Figure 4D). During normal ESC differentiation into neural progenitor cells (NPCs), H3K27me3 levels at these loci are reduced to permit gene activation, while mesoderm and endoderm marker genes retain the repressive H3K27me3 marks to ensure lineage restriction (Figure 4D). These findings raise the possibility that Pum1/2 depletion enhances PRC2 recruitment and H3K27me3 deposition at NE marker genes, suppressing their activation and contributing to reduced NE specification in differentiating mESCs.

## DISCUSSION

ESCs undergo extensive reprogramming of gene expression networks as they transition from the naïve pluripotent state to lineage-committed progenitors. Elucidating the molecular mechanisms governing this process has been a central topic for stem cell biology and is essential for applications such as organoid development and regenerative medicine. Here, we demonstrate that Pum1/2-mediated post-transcriptional regulation, though less explored than pluripotency and lineage-specific TFs, constitutes a critical layer of control over ESC fate decisions. Through systematic, high-temporal-resolution analyses, we uncover a two-phase cascade through which Pum1/2 reprogram gene expression during ESC differentiation.

### Identification of direct Pum1/2 targets and a biphasic regulatory hierarchy

Our optimized eCLIP enabled high-confidence identification of over 2,000 direct target mRNAs of Pum1/2 in mESCs, with precise mapping of their binding sites. Meanwhile, leveraging the fast-acting PROTAC technology, we achieved temporally acute Pum1/2 depletion in mESCs followed by time-course RNA-seq. This allowed us to capture over 100 mRNAs that were immediately stabilized within 4-10 hours of Pum1/2 depletion, most of which were direct Pum1/2 targets, defining the first regulatory phase dominated by the direct effect that down-regulates target mRNAs. From 10 to 72 hours, many additional mRNAs were regulated, either positively or negatively. Many of these mRNAs were not directly bound by Pum1/2, marking the onset of a second regulatory phase. While this phase still involves continuous regulation of direct Pum1/2 targets (Figure 3C), it is mostly characterized by indirect regulation mediated through early direct targets, with potential reinforcement by subsequent downstream regulators.

### Primary action mode of Pum1/2: negative control of mRNA stability

Combined RNA-seq and DIA-MS proteomics revealed that the majority of proteins upregulated upon Pum1/2 depletion corresponded to mRNAs with concordant increases in abundance. This finding indicates that Pum1 and Pum2 primarily promote target RNA degradation in mESCs, while a smaller subset of targets, such as *Cdkn1a*, may also be subject to translational repression.

Importantly, unlike previous reports^16,25^, we found no evidence that Pum1/2 stabilize their target mRNAs or enhance translation. If such effects were present, depletion of Pum1/2 would be expected to reduce the RNA or proteins levels of their targets. However, up to R6/D6 – time points at which direct effects of Pum1/2 loss predominate – all differentially expressed Pum1/2 targets were upregulated at both the RNA and protein levels. Downregulation of Pum1/2 target RNAs emerged only after R24/D24, when widespread changes in non-target genes were also observed. Therefore, this delayed downregulation likely reflects indirect responses rather than direct Pum1/2-mediated RNA stabilization or translational activation – a possibility that must be excluded before attributing such functions to Pum proteins. We suggest that previous conclusions of Pum1/2-mediated stabilization likely stemmed from technical limitations: conventional RNAi or gene knockout caused Pum1/2 depletion well before ESCs were sampled for transcriptome and translation analyses. This prolonged loss of Pum1/2 can cause potentially indirect effects on Pum1/2-target mRNAs.

Another key insight from our study is that Pum1/2 often act as fine-tuners of target RNA stability, reducing steady-state levels by less than two-fold. Nevertheless, even these modest effects can modulate the expression of over 1,000 genes and substantially influence ESC fate decisions, likely owing to the large number of direct regulatory targets (>100) and the potent downstream regulatory effects of specific targets, such as *Gata6*^15^.

### Pum1/2 are important for ESC pluripotency and lineage balance

Previous studies of Pum1/2 functions were focused on a narrow set of pluripotency and lineage markers. In contrast, by performing time-resolved RNA-seq over a 72-hour period following LIF/2i withdrawal, we delineated the transcriptional trajectories underlying lineage specification events in Pum1/2-depleted mESCs. This systematic approach uncovers a previously unrecognized role for Pum proteins in balancing NE and PGC fate choices during early embryogenesis. This may be partly achieved by fine-tuning mRNA stability of PRC2 components, such as *Suz12*. By promoting mRNA degradation of these key epigenetic regulators, Pum1/2 likely facilitate dynamic reprogramming of chromatin modifications and transcriptional networks, a process essential for cell fate transitions. These findings underscore the importance of post-transcriptional regulation in coordinating developmental gene expression programs.

## METHODS

### mES cell culture and differentiation

The naïve mouse ESCs were cultured on 0.1% gelatin-coated plates in the serum/LIF/2i medium as previously described^36^. Briefly, cells were grown in DMEM High Glucose (Gibco, 11965-092) supplemented with 15% fetal bovine serum (FBS, Gibco, 26140-079), 2 mM GlutaMAX (Gibco, 35050-061), 1 mM sodium pyruvate (Gibco, 11360-070), 1x minimum essential medium nonessential amino acids (MEM NEAA, Gibco, 11140-050), 0.1 mM 2-mercaptoethanol (Sigma, M3148), 1000 U/mL Leukemia inhibitory factor (LIF, Sigma, ESG1107), 1 µM PD0325901 (Sigma, 444968), and 3 µM CHIR99021 (Sigma, 361571). Medium change was performed every 1-2 days.

mES cells were passaged every 3-4 days by aspirating the old medium, dissociating with TrypLE for 3 min at 37°C, and then rinsing and further dissociation in serum/LIF/2i medium by pipetting. Cells were pelleted by centrifugation at 200-500 g for 5 min, resuspended in culture medium, and seeded 5,000-10,000 cells/cm^2^ in tissue-culture plates. Cell counting was performed using hemocytometer.

For induction of spontaneous differentiation, ESCs were grown in ESC differentiation medium: DMEM high glucose supplemented with 15% FBS, 2 mM GlutaMax, 1 mM sodium pyruvate, 1x MEM NEAA and 0.1 mM 2-mercaptoethanol, in the absence of LIF or 2 inhibitors (MEKi, GSK3i).

### mES cell electroporation

Cells were electroporated using the Nucleofector Kit for mouse embryonic stem cells (Lonza, VPH-1001) with program A-030. For each electroporation, 2-3x10^6^ cells were used. For CRISPR-mediated knock-in experiments, 2 µg each of PX458/459 plasmid (Cas9 and sgRNA) and 7 µg HDR repair template were used. For transient transfection experiments, 2 µg of plasmid was used for each electroporation. Cells post electroporation were seeded to 0.1% gelatin-coated T75 flasks in serum/LIF/2i medium and cultured for 2-3 days before selection with antibiotics.

### Generation of ESC knock-in cell lines

ESCs were co-transfected with one or two Cas9-sgRNA plasmids (depending on the number of sgRNAs needed) and the HDR donor-vector through electroporation, and selected 48-72h later with 100 µg/mL hygromycin B (Gibco, 10687-010), 150 µg/mL G418 (Gibco, 10131-035), or 4 µg/mL blasticidin S (Gibco, A1113903). After six to seven days post-electroporation, the cells were sorted by flow cytometry (BD FACS Aria) and single cells were seeded to each well of a 0.1% gelatin-coated 96-well plate. Around 10 days after sorting, clonal ESCs were expanded in 48-well plates. PCR genotyping was performed to screen for homozygous or heterozygous knock-ins, and all homozygous knock-in cell lines were confirmed by Sanger sequencing. Western blot was performed to confirm expression of fusion proteins after CRISPR-mediated knock-ins.

### Construction of Cas9-sgRNA vectors

The PX458 (addgene #48138) or PX459 (addgene #62988) vector was linearized with BbsI-HF digestion and gel purified. A pair of DNA oligos for each targeting site were phosphorylated, annealed, and ligated to the linearized PX458 or PX459 as previously described^37^.

### Construction of HDR donor-vectors

The HDR donor-vectors were assembled from three to six DNA fragments including a linearized pBlueScript backbone using Gibson Assembly Master Mix (NEB, E2611L). The DNA fragments were PCR amplified from ESC genomic DNA (homology arms for HDR) or other DNA plasmids. The pBlueScript backbone was linearized by digestion with NotI-HF and KpnI-HF. All DNA fragments were gel purified before the Gibson assembly reactions.

### Western blot

Cells were lysed in RIPA buffer (50 mM Tris-HCl pH 8.0, 150 mM NaCl, 1.0% Triton X-100, 0.1% SDS, 1 mM EDTA, 0.5% sodium deoxycholate) supplemented with cOmplete Mini EDTA-free protease inhibitor cocktail (Roche, #11836170001) and PhosSTOP (Roche, #4906845001). Lysates were denatured in 1x Laemmli buffer at 70C for 10 min. Proteins were separated by SDS-PAGE using acrylamide gels (mini-protean) and transferred onto 0.45 µm PVDF membranes at 16V for 45-60 min using Trans-Blot SD Semi-Dry Transfer Cell. The PVDF membranes were blocked in 5% non-fat milk in PBST (0.1% Tween-20 in PBS) for 30-60 min at room temperature (RT), and then incubated with primary antibodies for 1-2 h at RT. After three washes with PBST, the PVDF membranes were incubated with HRP-conjugated secondary antibodies at RT for 1 h. The signal was developed with SuperSignal^TM^ West Pico PLUS Chemiluminescent Substrate (Thermo Scientific, 34580).

### RNA-seq and data analysis

Total RNA was extracted from ESCs with RNeasy Plus Mini Kit (Qiagen, 74134) or Quick-RNA MagBead kit (Zymo Research, R2132). For Quick-RNA MagBead kit, genomic DNA was removed by DNase I treatment for 10 min at 37°C. 1 µg of total RNA was used for poly(A)-selected library preparation with KAPA mRNA HyperPrep Kit (Roche) following manufacturer’s protocols. RNA-seq libraries were sequenced on the Illumina NovaSeq X Plus platform (paired-end 2x100 bp).

Sequencing quality was assessed using FastQC. Low-quality reads and adapter sequences were removed with Cutadapt, and the trimmed reads were aligned to the mouse reference genome (GRCm39) using STAR. Read counts were generated with featureCounts, and differential gene expression analysis was performed with DESeq2 in R.

### eCLIP library preparation

UV-crosslinking was performed as previously described^38^. Briefly, mESCs (LIF+2i) cultured in 10cm plates were trypsinized, washed, and diluted in ice-cold 1X DPBS to a density of ∼10 million cells per mL, and 6 mL of single-cell suspension was added to each 10cm dish for protein-RNA crosslinking. Cells were irradiated on ice with 500 mJ/cm^2^ of UV at 254 nm in a Stratalinker 2400. Cells were then transferred to 50 mL tubes, washed once with 1X DPBS supplemented with 2% BSA, aliquoted in 1.5 mL tubes (∼25 million cells each), and pelleted by centrifugation at 500 g for 5 min. Cell pellets were snap-frozen on dry ice and stored at -80°C until further use.

eCLIP was performed as previously described^39^, with some modifications to streamline the workflow (see Table S4 for oligonucleotides used). For each replicate, ∼25 million UV-crosslinked mESCs were lysed on ice for 15 min in 1 mL ice-cold iCLIP lysis buffer (50 mM Tris-HCl pH 7.5, 100 mM NaCl, 1% NP-40, 0.1% SDS, 0.5% sodium deoxycholate) supplemented with cOmplete Mini EDTA-free protease inhibitor cocktail (Roche, #11836170001) and 12.5 µL Murine RNase Inhibitor (NEB, M0314L). Cell lysates were then sonicated in Bioruptor (Diagenode, UCD-200) for 5 min at 4°C (“Mid” setting, pulses 30 sec on, 30 sec off) and cleared by centrifugation for 15 min at 15,000 g, 4°C. The supernatant was transferred to new tubes containing pre-coupled antibody-beads (50 µL or 1.5 mg Dynabead Protein G + 10 µg antibodies) and left rotating overnight (∼ 16 h) at 4°C to capture RBP-RNA complexes on beads.

After overnight immunoprecipitation, RNase I (100 U/µL, Invitrogen, AM2294) was added to the cell lysate-bead suspension (∼1 mL) at a final concentration of 200 U/mL (1:500 dilution) along with 10 µL Turbo DNase (Invitrogen, AM2238), and samples were incubated at 37°C for 30 min in Thermomixer (1200 rpm shaking) for partial RNA fragmentation to approximately 30∼150 nt in length. RNase reaction was quenched on ice with addition of 12 µL SUPERase·In^TM^ RNase inhibitor (20 U/µL, Invitrogen, AM2696). For each replicate, 21 µL (2%) of cell lysate-bead suspension was saved at 4°C for library preparation of the pre-IP input controls.

Protein-RNA complexes captured by the magnetic beads were washed 3 times with high-salt wash buffer (50 mM Tris-HCl pH 7.5, 1 M NaCl, 1 mM EDTA, 1% NP-40, 0.1% SDS, 0.5% sodium deoxycholate), once with wash buffer (20 mM Tris-HCl pH 7.5, 10 mM MgCl_2_, 0.2% Tween-20), and twice with 1X FastAP buffer (10 mM Tris-HCl pH 7.5, 100 mM KCl, 5 mM MgCl_2_, 0.2% Triton X-100). RNAs were then dephosphorylated on-bead with FastAP and T4 PNK, and ligated with RNA_N5_biotin adapter by T4 RNA ligase 1 (NEB, M0437M) at 25°C overnight (∼16 h) in Thermomixer (1200 rpm shaking). After ligation, RBP-RNA complexes were washed, denatured, run on NuPAGE 4-12% Bis-Tris Gel (Invitrogen, NP0335BOX), and transferred to nitrocellulose membranes (0.22 µm) at 30V overnight (∼16 h) in an XCell II blot module. Regions above the target protein size (Pum1: 150-300 kDa; Pum2: 125-250 kDa) were excised from the nitrocellulose membrane, treated with Proteinase K (NEB, P8107S) to release RNA, and RNA was purified with RNA Clean & Concentrator-5 (Zymo, R1016).

Without gel electrophoresis and membrane transfer, pre-IP input samples (bead suspension) were directly treated with Proteinase K. RNA was then purified, dephosphorylated, and 3’-ligated to RNA_N5_biotin adapter.

Purified 3’-ligated RNAs (both CLIP and INPUT) were reverse transcribed with SuperScript III (Invitrogen, 18080044) using AR21 primer, followed by treatment with ExoSAP-IT (Applied Biosystems, 78200.200.UL) to remove excess primers and NaOH (final conc. 100 mM) to remove RNA. The purified cDNAs were then 3’-ligated to the rand10_3Tr3 adapter with T4 RNA ligase 1 at 25°C overnight (∼16 h) in Thermomixer (1200 rpm shaking). After clean-up with Dynabeads MyOne Silane (Invitrogen, 37002D), 1 µL of the 1:10 diluted cDNA was used for qPCR quantification (P5 and P7 Solexa primers) and 10 µL of undiluted cDNA (total elution 25 µL) was PCR amplified with Q5 Hot Start High-Fidelity DNA Polymerase (NEB, M0494S) and D50x/D70x primers (3 cycles less than the Ct in qPCR). The PCR reactions were treated with ExoSAP-IT to remove excess primers and cleaned up using 1.2X Ampure XP beads (size-selection for DNA fragments >175 bp). The resulting eCLIP libraries were sequenced on the Illumina NovaSeq 6000 platform in paired-end 2x100 bp mode.

### eCLIP-seq data processing

Analysis of eCLIP data was performed as previously described^39^. Paired-end sequencing reads were quality-checked by FastQC and trimmed to remove adapter sequences by Cutadapt (two rounds of trimming). The unique molecule identifiers (UMIs) – 5 or 10 random bases at the 5’ ends of R1 and R2, respectively – were trimmed and appended to the read names using a custom script. The trimmed read pairs were then aligned to the mouse reference genome (GRCm39) using STAR, and only the uniquely mapped reads were used for further analysis. PCR duplicate reads sharing the same R1 start position, R2 start position, and UMIs were removed using a custom script, and then R2 of the read pairs were kept for further analysis. Peak calling was performed using CLIPper^40^ (downloaded from https://github.com/YeoLab/clipper.git), and normalization of peak signals against pre-IP input was performed using a custom script. Briefly, the numbers of CLIP and INPUT reads overlapping the CLIPper-called peaks were used to calculate fold enrichment (normalized by total read numbers in each data set), and the enrichment p-value was calculated by Yates’ Chi-Square test, or Fisher’s Exact Test when read number < 5 in CLIP or INPUT. Peaks with log_2_FC > 4.5 and p < 1x10^-10^ were considered significantly enriched over input, and the genes with at least one significant peak on their mRNA transcripts were identified as the target genes.

### Mass spectrometry-based proteomics: sample preparation and data processing

The lysis buffer containing 10 M urea containing the cOmplete™ protease inhibitor cocktail (Roche, #11697498001) was used for protein extraction on cell samples. The cell lysates in 2 mL tubes were completely lysed by sonication at 4°C for two cycle (1 min per cycle) using a VialTweeter device (Hielscher-Ultrasound Technology). The lysed samples were then centrifuged at 20,000 x g for 1 hour to remove insoluble material. Identical amounts of proteins were utilized for the subsequent trypsin digestion process. Reduction and alkylation were carried out using 10 mM Dithiothreitol (DTT) for 1 hour at 56°C, followed by 20 mM iodoacetamide (IAA) in darkness for 45 minutes at room temperature. The samples were then diluted with 100 mM NH_4_HCO_3_ and digested with trypsin (Promega) at a ratio of 1:20 (w/w) overnight at 37°C with shaking at 600 rpm. The purification of the digested peptides was performed using a C18 column (MacroSpin Columns, NEST Group INC). 1 µg of the peptide was utilized for total proteome analysis.

The samples were measured by DIA-MS as described previously^41–43^, on an Orbitrap Fusion Tribrid mass spectrometer (Thermo Scientific) coupled to a nanoelectrospray ion source (NanoFlex, Thermo Scientific) and an EASY-nLC 1200 system (Thermo Scientific, San Jose, CA). A 150 min or 120 min gradient was used for the data acquisition at the flow rate at 300 nL/min with the column temperature controlled at 60 °C using a column oven (PRSO-V1, Sonation GmbH, Biberach, Germany). DIA-MS consisted of one MS1 scan and 33 MS2 scans of variable isolated windows with 1 m/z overlapping between windows. The MS1 scan range was 350 – 1650 m/z and the MS1 resolution was 120,000 at m/z 200. The MS1 full scan AGC target value was set to be 2E6 and the maximum injection time was 100 ms. The MS2 resolution was set to 30,000 at m/z 200 with the MS2 scan range 200 – 1800 m/z and the normalized HCD collision energy was 28%. The MS2 AGC was set to be 1.5E6 and the maximum injection time was 50 ms. The default peptide charge state was set to 2. Both MS1 and MS2 spectra were recorded in profile mode. DIA-MS data was analyzed using Spectronaut v19^44–46^ with directDIA algorithm by searching against the SwissProt downloaded mouse fasta file (Sep 2024). The oxidation at methionine was set as variable modification, whereas carbamidomethylation at cysteine was set as fixed modification. Both peptide and protein FDR cutoffs (Qvalue) were controlled below 1% and the resulting quantitative data matrix were exported from Spectronaut. All the other settings in Spectronaut were kept as Default.

### ChIP-seq data analysis

Published ChIP-sea data was download using SRA toolkit. Basic quality control was performed on the FASTQ files using FastQC. Without trimming adapters, the sequencing reads were aligned to the reference mouse genome (GRCm39) using Bowtie2.

The resulting SAM files were converted to BAM files using SAMtools. The PCR duplicates were removed using a customized script, and the de-duplicate BAM files were sorted and indexed. BAM files were converted to BigWig files using the bamCoverage function of DeepTools. The IGV genome browser was used to visualize the signals of ChIP-seq. DeepTools was also used to generate the read coverage heatmaps.

## AVAILABILITY OF DATA AND MATERIALS

The eCLIP and RNA-seq data generated in this study are available in the GEO database with the accession numbers GSE310693 and GSE310694. The mass spectrometry data have been deposited to the ProteomeXchange Consortium via the PRIDE partner repository: PXD069838. Other data are available from the corresponding author upon reasonable request.

## COMPETING INTERESTS

No competing interest is declared.

## AUTHOR CONTRIBUTIONS STATEMENT

Y.H. and H.L. conceived the project. Y.H. conducted most of the experiments and bioinformatic analyses. W.L. and Y.L. conducted mass spectrometry experiments. H.R. helped validate the efficacy of PROTAC systems. Y.H. and H.L. wrote the manuscript.

## ACKNOWLEDGEMENTS

We thank Jitong Cai for bioinformatic support, Salil Garg for providing the Nanog-mCerulean3 mESCs, and members of Haifan Lin’s lab for insightful discussions. We also thank the MS & Proteomics Resource at Yale University for conducting mass spectrometry and related experiments.

---

**Figure S1.**
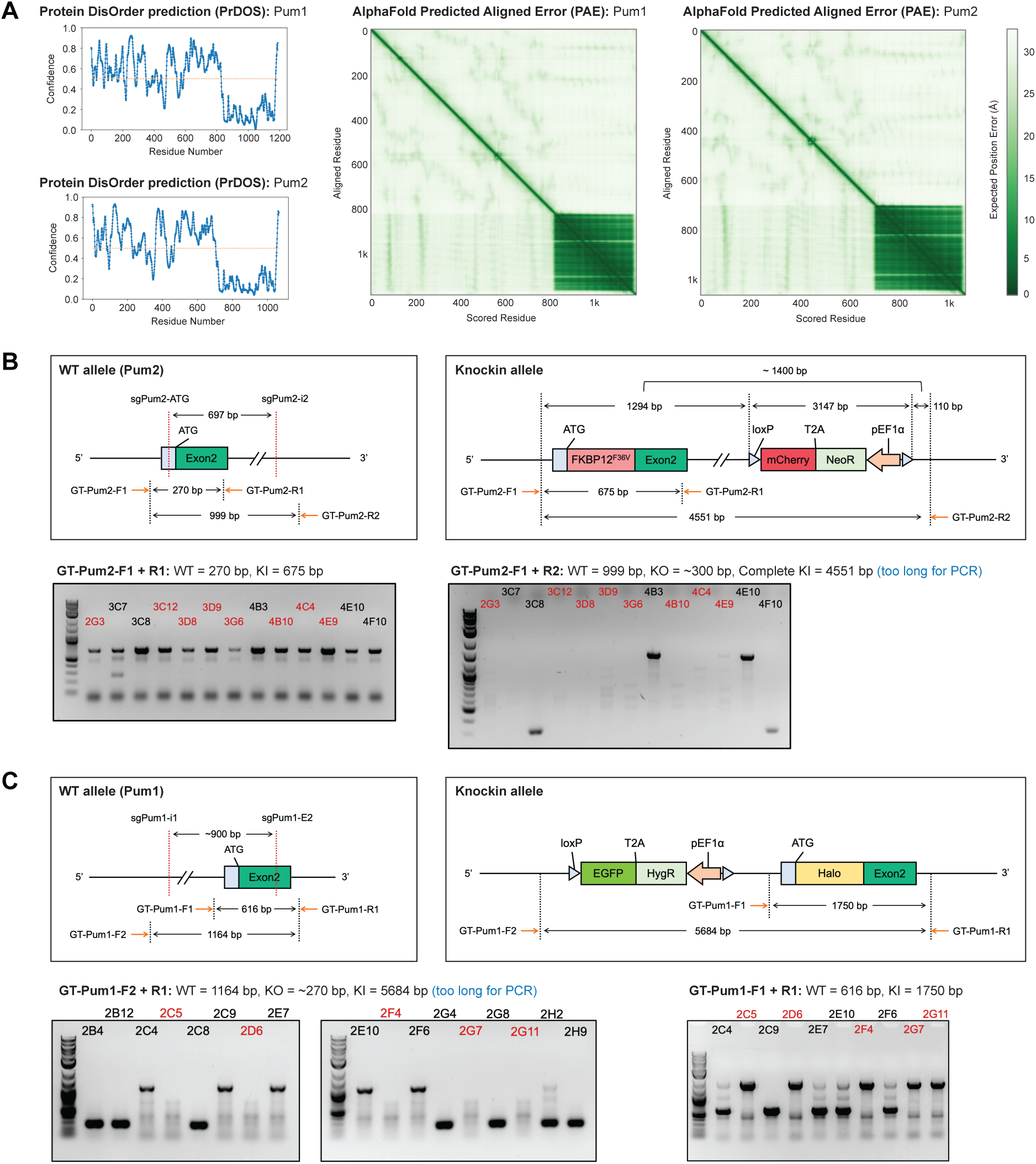
Related to Figure 1 Generation of *Pum1*^Halo/Halo^; *Pum2*^dTAG/dTAG^ mESCs. (A) PrDOS and AlphaFold predictions of intrinsically disordered regions (IDRs) in Pum1 and Pum2 proteins. In the PrDOS plots, the y-axis represents the prediction confidence score, with higher values indicating a greater likelihood of intrinsic disorder. (B) Validation of FKBP12^F36V^-Pum2 CRISPR knock-in clones by PCR on genomic DNA. Clones highlighted in red represent potential homozygous knock-ins. (C) Validation of Halo-Pum1 CRISPR knock-in clones by PCR on genomic DNA. Clones highlighted in red represent potential homozygous knock-ins.

**Figure S2.**
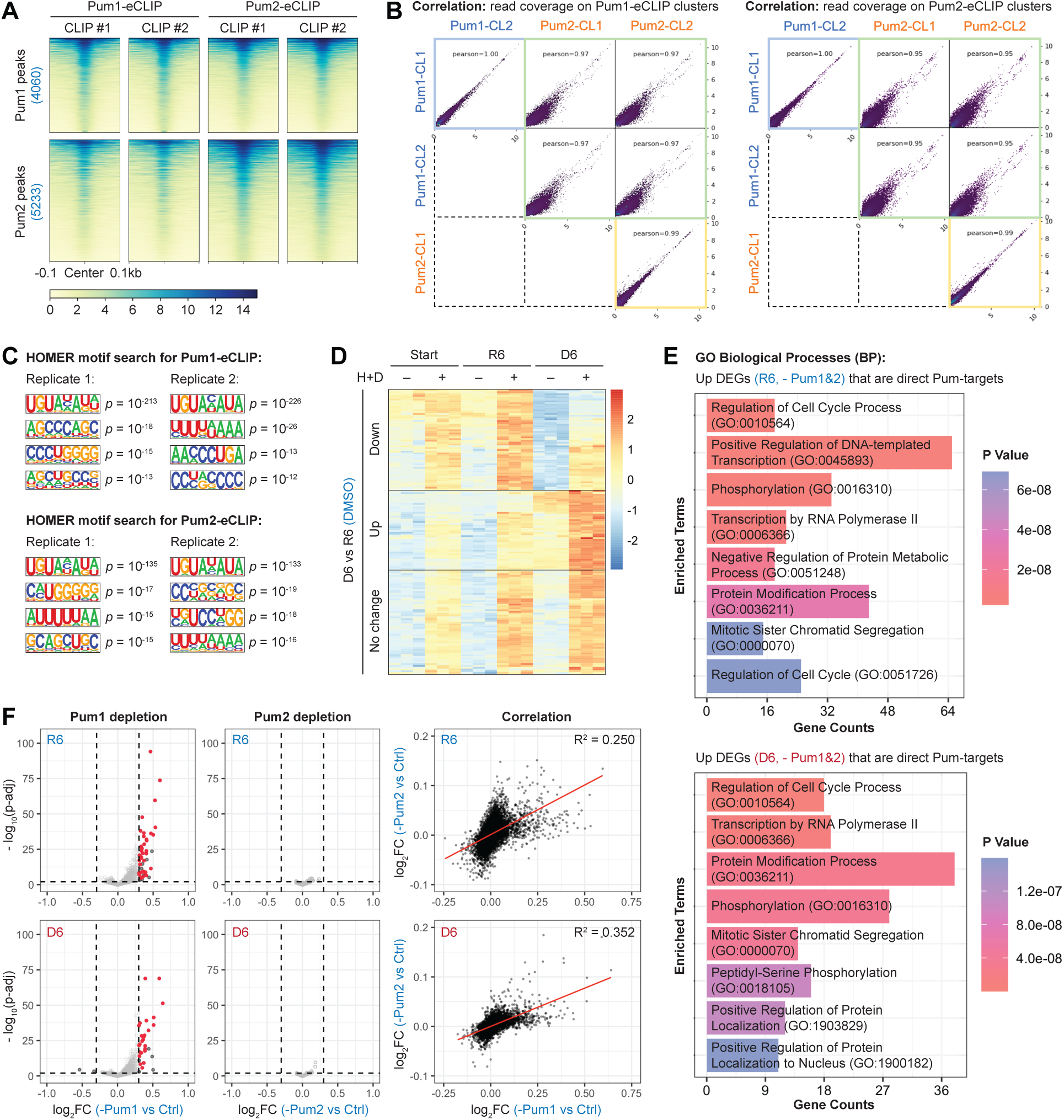
Related to Figure 2 Pum proteins predominantly act by destabilizing their target RNAs. (A) Heatmaps showing coverage of Pum1- and Pum2-eCLIP reads (two biological replicates) within significantly enriched Pum1- and Pum2-binding clusters. (B) Scatter plots showing correlations among read coverage of Pum1- and Pum2-eCLIP samples on Pum1- and Pum2-binding clusters. (C) HOMER *de novo* motif search results of Pum1- and Pum2-eCLIP, showing the top 4 motifs for each replicate. Background sequences were generated by randomly shuffling the bases within the significantly enriched Pum1/2-binding peaks to match the base content of the peak sequences. (D) Heatmap showing DESeq2-normalized counts of the DEGs (log_2_FC > 0.3, adjusted p < 0.01) in DMSO-treated control and dual Pum1/2-depleted mESCs at Start, R6, and D6. Colors represent the z-scores (normalized to row means). (E) Gene ontology (GO) analysis (biological processes, BP) for the DEGs (log_2_FC > 0, adjusted p < 0.01) in dual Pum1/2-depleted mESCs at R6 and D6. (F) Left and middle charts: volcano plots showing differential regulation of RNAs in Pum1- or Pum2-depleted mESCs at R6 and D6. Significance cutoffs: |log_2_FC| > 0.3, adjusted p-value < 0.01. The direct Pum1/2-target mRNAs are highlighted in red. Right charts: scatter plots showing log_2_FC correlation between Pum1- and Pum2-depleted mESCs at R6 and D6.

**Figure S3.**
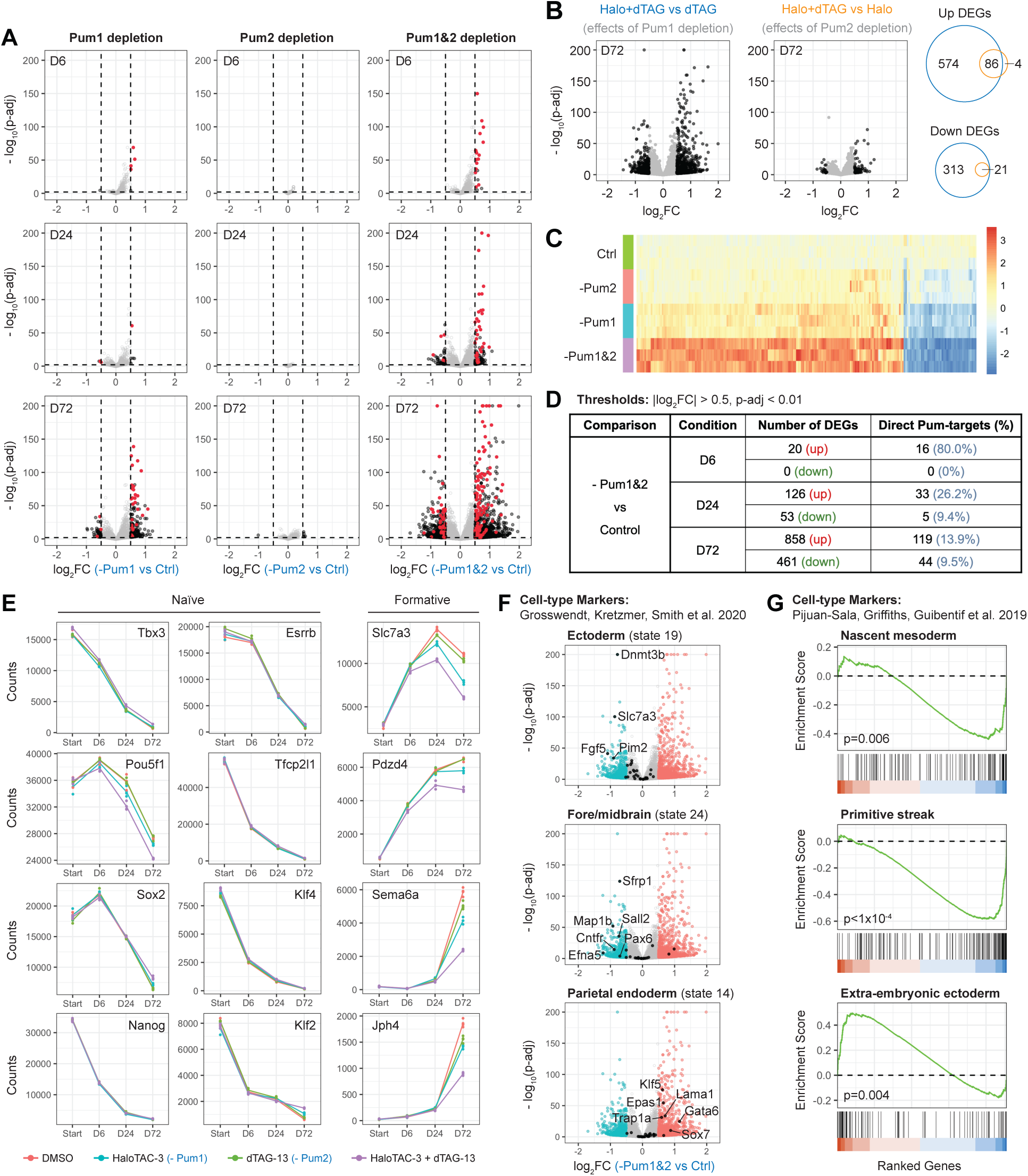
Related to Figure 3 Depletion of Pum1 and Pum2 disrupts cell fate decisions of mESCs. (A) Volcano plots showing differential regulation of RNAs in Pum1- and/or Pum2-depleted mESCs at the indicated time points. Significance cutoffs: |log_2_FC| > 0.5, adjusted p-value < 0.01. The direct Pum1/2-target mRNAs are highlighted in red. (B) Volcano plots of DEGs at D72, comparing Pum1/2 double-depletion versus single-depletion. Venn plots show the overlap between the two sets of DEGs. (C) Heatmap showing DESeq2-normalized counts of the DEGs identified in (B) at D72. Colors represent the z-scores (normalized to mean of control for each column/gene). (D) Summary of the DEG numbers in Pum1/2-double-depleted mESCs at the indicated time points, and the numbers and percentages of direct Pum1/2-target RNAs in the DEGs. (E) Trend plots showing the mRNA levels (DESeq2-normalized counts) of selected naïve and formative pluripotency markers during mESC differentiation (n = 3). (F) Volcano plots showing DEGs at D72 (dual Pum1/2-depletion vs. control), with marker genes for ectoderm, fore/midbrain, or parietal endoderm highlighted in black. (G) GSEA for nascent mesoderm, primitive streak, or extra-embryonic ectoderm markers at D72 (dual Pum1/2-depletion vs. control).

**Figure S4.**
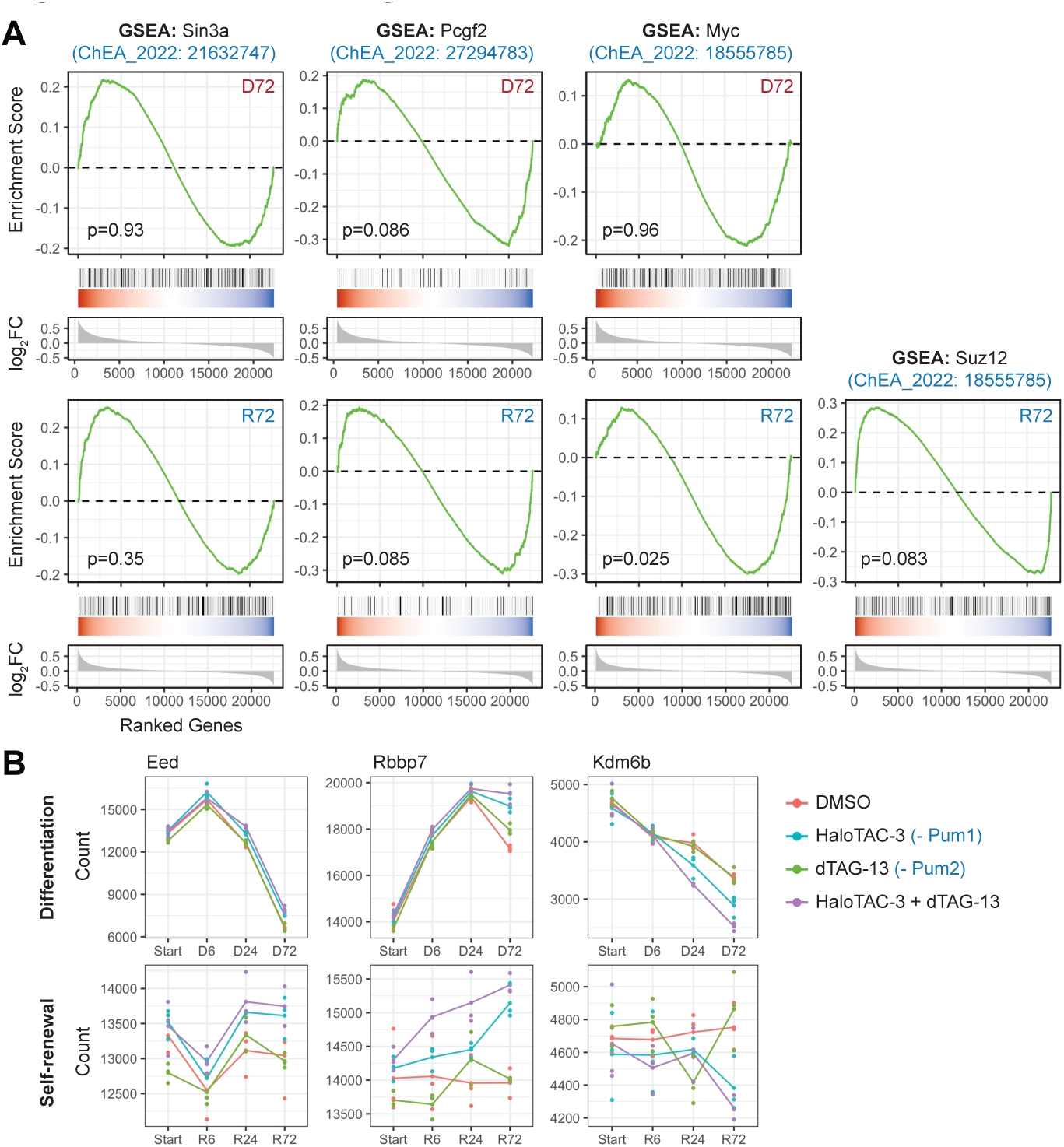
Related to Figure 4 Pum1 and Pum2 regulate PRC2 and H3K27me3 during mESC differentiation. (A) GSEA for the indicated TF-target gene sets at D72 and R72 (Pum1/2 double-depletion vs. control). (B) Trend plots showing mRNA levels (DESeq2-normalized counts) of PRC2 components and H3K27me3 demethylase genes during mESC differentiation and self-renewal (n = 3).

## Notes

### Competing Interest Statement

The authors have declared no competing interest.

## REFERENCES

1. Hemberger, M., Dean, W., and Reik, W. (2009). Epigenetic dynamics of stem cells and cell lineage commitment: digging Waddington’s canal. Nat Rev Mol Cell Biol 10, 526–537. 10.1038/nrm2727.

2. Meissner, A. (2010). Epigenetic modifications in pluripotent and differentiated cells. Nat Biotechnol 28, 1079–1088. 10.1038/nbt.1684.

3. Yeo, J.C., and Ng, H.H. (2013). The transcriptional regulation of pluripotency. Cell Res 23, 20–32. 10.1038/cr.2012.172.

4. Kravchenko, P., and Tachibana, K. (2025). Rise and SINE: roles of transcription factors and retrotransposons in zygotic genome activation. Nat Rev Mol Cell Biol 26, 68–79. 10.1038/s41580-024-00772-6.

5. Lu, R., Markowetz, F., Unwin, R.D., Leek, J.T., Airoldi, E.M., MacArthur, B.D., Lachmann, A., Rozov, R., Ma’ayan, A., Boyer, L.A., et al. (2009). Systems-level dynamic analyses of fate change in murine embryonic stem cells. Nature 462, 358–362. 10.1038/nature08575.

6. Chen, Q., and Hu, G. (2017). Post-transcriptional regulation of the pluripotent state. Curr Opin Genet Dev 46, 15–23. 10.1016/j.gde.2017.06.010.

7. Mohammed, H., Hernando-Herraez, I., Savino, A., Scialdone, A., Macaulay, I., Mulas, C., Chandra, T., Voet, T., Dean, W., Nichols, J., et al. (2017). Single-Cell Landscape of Transcriptional Heterogeneity and Cell Fate Decisions during Mouse Early Gastrulation. Cell Rep 20, 1215–1228. 10.1016/j.celrep.2017.07.009.

8. Pijuan-Sala, B., Griffiths, J.A., Guibentif, C., Hiscock, T.W., Jawaid, W., Calero-Nieto, F.J., Mulas, C., Ibarra-Soria, X., Tyser, R.C.V., Ho, D.L.L., et al. (2019). A single-cell molecular map of mouse gastrulation and early organogenesis. Nature 566, 490–495. 10.1038/s41586-019-0933-9.

9. Cheng, S., Pei, Y., He, L., Peng, G., Reinius, B., Tam, P.P.L., Jing, N., and Deng, Q. (2019). Single-Cell RNA-Seq Reveals Cellular Heterogeneity of Pluripotency Transition and X Chromosome Dynamics during Early Mouse Development. Cell Rep 26, 2593–2607 e2593. 10.1016/j.celrep.2019.02.031.

10. Mittnenzweig, M., Mayshar, Y., Cheng, S., Ben-Yair, R., Hadas, R., Rais, Y., Chomsky, E., Reines, N., Uzonyi, A., Lumerman, L., et al. (2021). A single-embryo, single-cell time-resolved model for mouse gastrulation. Cell 184, 2825–2842 e2822. 10.1016/j.cell.2021.04.004.

11. Qiu, C., Cao, J., Martin, B.K., Li, T., Welsh, I.C., Srivatsan, S., Huang, X., Calderon, D., Noble, W.S., Disteche, C.M., et al. (2022). Systematic reconstruction of cellular trajectories across mouse embryogenesis. Nat Genet 54, 328–341. 10.1038/s41588-022-01018-x.

12. Thomson, M., Liu, S.J., Zou, L.N., Smith, Z., Meissner, A., and Ramanathan, S. (2011). Pluripotency factors in embryonic stem cells regulate differentiation into germ layers. Cell 145, 875–889. 10.1016/j.cell.2011.05.017.

13. Grosswendt, S., Kretzmer, H., Smith, Z.D., Kumar, A.S., Hetzel, S., Wittler, L., Klages, S., Timmermann, B., Mukherji, S., and Meissner, A. (2020). Epigenetic regulator function through mouse gastrulation. Nature 584, 102–108. 10.1038/s41586-020-2552-x.

14. Leeb, M., Dietmann, S., Paramor, M., Niwa, H., and Smith, A. (2014). Genetic exploration of the exit from self-renewal using haploid embryonic stem cells. Cell Stem Cell 14, 385–393. 10.1016/j.stem.2013.12.008.

15. Lin, K., Zhang, S., Shi, Q., Zhu, M., Gao, L., Xia, W., Geng, B., Zheng, Z., and Xu, E.Y. (2018). Essential requirement of mammalian Pumilio family in embryonic development. Mol Biol Cell 29, 2922–2932. 10.1091/mbc.E18-06-0369.

16. Uyhazi, K.E., Yang, Y., Liu, N., Qi, H., Huang, X.A., Mak, W., Weatherbee, S.D., de Prisco, N., Gennarino, V.A., Song, X., and Lin, H. (2020). Pumilio proteins utilize distinct regulatory mechanisms to achieve complementary functions required for pluripotency and embryogenesis. Proc Natl Acad Sci U S A 117, 7851–7862. 10.1073/pnas.1916471117.

17. Zhang, M., Chen, D., Xia, J., Han, W., Cui, X., Neuenkirchen, N., Hermes, G., Sestan, N., and Lin, H. (2017). Post-transcriptional regulation of mouse neurogenesis by Pumilio proteins. Genes Dev 31, 1354–1369. 10.1101/gad.298752.117.

18. Buckley, D.L., Raina, K., Darricarrere, N., Hines, J., Gustafson, J.L., Smith, I.E., Miah, A.H., Harling, J.D., and Crews, C.M. (2015). HaloPROTACS: Use of Small Molecule PROTACs to Induce Degradation of HaloTag Fusion Proteins. ACS Chem Biol 10, 1831–1837. 10.1021/acschembio.5b00442.

19. Nabet, B., Roberts, J.M., Buckley, D.L., Paulk, J., Dastjerdi, S., Yang, A., Leggett, A.L., Erb, M.A., Lawlor, M.A., Souza, A., et al. (2018). The dTAG system for immediate and target-specific protein degradation. Nat Chem Biol 14, 431–441. 10.1038/s41589-018-0021-8.

20. Ishida, T., and Kinoshita, K. (2007). PrDOS: prediction of disordered protein regions from amino acid sequence. Nucleic Acids Res 35, W460–464. 10.1093/nar/gkm363.

21. Fleming, J., Magana, P., Nair, S., Tsenkov, M., Bertoni, D., Pidruchna, I., Lima Afonso, M.Q., Midlik, A., Paramval, U., Zidek, A., et al. (2025). AlphaFold Protein Structure Database and 3D-Beacons: New Data and Capabilities. J Mol Biol 437, 168967. 10.1016/j.jmb.2025.168967.

22. Jumper, J., Evans, R., Pritzel, A., Green, T., Figurnov, M., Ronneberger, O., Tunyasuvunakool, K., Bates, R., Zidek, A., Potapenko, A., et al. (2021). Highly accurate protein structure prediction with AlphaFold. Nature 596, 583–589. 10.1038/s41586-021-03819-2.

23. Kedde, M., van Kouwenhove, M., Zwart, W., Oude Vrielink, J.A., Elkon, R., and Agami, R. (2010). A Pumilio-induced RNA structure switch in p27-3’ UTR controls miR-221 and miR-222 accessibility. Nat Cell Biol 12, 1014–1020. 10.1038/ncb2105.

24. Etersque, J.M., Lee, I.K., Sharma, N., Xu, K., Ruff, A., Northrup, J.D., Sarkar, S., Nguyen, T., Lauman, R., Burslem, G.M., and Sellmyer, M.A. (2023). Regulation of eDHFR-tagged proteins with trimethoprim PROTACs. Nat Commun 14, 7071. 10.1038/s41467-023-42820-3.

25. Bohn, J.A., Van Etten, J.L., Schagat, T.L., Bowman, B.M., McEachin, R.C., Freddolino, P.L., and Goldstrohm, A.C. (2018). Identification of diverse target RNAs that are functionally regulated by human Pumilio proteins. Nucleic Acids Res 46, 362–386. 10.1093/nar/gkx1120.

26. Goldstrohm, A.C., Hall, T.M.T., and McKenney, K.M. (2018). Post-transcriptional Regulatory Functions of Mammalian Pumilio Proteins. Trends Genet 34, 972–990. 10.1016/j.tig.2018.09.006.

27. Wang, X., Xiang, Y., Yu, Y., Wang, R., Zhang, Y., Xu, Q., Sun, H., Zhao, Z.A., Jiang, X., Wang, X., et al. (2021). Formative pluripotent stem cells show features of epiblast cells poised for gastrulation. Cell Res 31, 526–541. 10.1038/s41422-021-00477-x.

28. Smith, A. (2017). Formative pluripotency: the executive phase in a developmental continuum. Development 144, 365–373. 10.1242/dev.142679.

29. Diamant, I., Clarke, D.J.B., Evangelista, J.E., Lingam, N., and Ma’ayan, A. (2025). Harmonizome 3.0: integrated knowledge about genes and proteins from diverse multi-omics resources. Nucleic Acids Res 53, D1016–D1028. 10.1093/nar/gkae1080.

30. Rouillard, A.D., Gundersen, G.W., Fernandez, N.F., Wang, Z., Monteiro, C.D., McDermott, M.G., and Ma’ayan, A. (2016). The harmonizome: a collection of processed datasets gathered to serve and mine knowledge about genes and proteins. Database (Oxford) 2016. 10.1093/database/baw100.

31. Shen, X., Liu, Y., Hsu, Y.J., Fujiwara, Y., Kim, J., Mao, X., Yuan, G.C., and Orkin, S.H. (2008). EZH1 mediates methylation on histone H3 lysine 27 and complements EZH2 in maintaining stem cell identity and executing pluripotency. Mol Cell 32, 491–502. 10.1016/j.molcel.2008.10.016.

32. Chamberlain, S.J., Yee, D., and Magnuson, T. (2008). Polycomb repressive complex 2 is dispensable for maintenance of embryonic stem cell pluripotency. Stem Cells 26, 1496–1505. 10.1634/stemcells.2008-0102.

33. Leeb, M., Pasini, D., Novatchkova, M., Jaritz, M., Helin, K., and Wutz, A. (2010). Polycomb complexes act redundantly to repress genomic repeats and genes. Genes Dev 24, 265–276. 10.1101/gad.544410.

34. Pasini, D., Cloos, P.A., Walfridsson, J., Olsson, L., Bukowski, J.P., Johansen, J.V., Bak, M., Tommerup, N., Rappsilber, J., and Helin, K. (2010). JARID2 regulates binding of the Polycomb repressive complex 2 to target genes in ES cells. Nature 464, 306–310. 10.1038/nature08788.

35. Kloet, S.L., Karemaker, I.D., van Voorthuijsen, L., Lindeboom, R.G.H., Baltissen, M.P., Edupuganti, R.R., Poramba-Liyanage, D.W., Jansen, P., and Vermeulen, M. (2018). NuRD-interacting protein ZFP296 regulates genome-wide NuRD localization and differentiation of mouse embryonic stem cells. Nat Commun 9, 4588. 10.1038/s41467-018-07063-7.

36. Czechanski, A., Byers, C., Greenstein, I., Schrode, N., Donahue, L.R., Hadjantonakis, A.K., and Reinholdt, L.G. (2014). Derivation and characterization of mouse embryonic stem cells from permissive and nonpermissive strains. Nat Protoc 9, 559–574. 10.1038/nprot.2014.030.

37. Ran, F.A., Hsu, P.D., Wright, J., Agarwala, V., Scott, D.A., and Zhang, F. (2013). Genome engineering using the CRISPR-Cas9 system. Nat Protoc 8, 2281–2308. 10.1038/nprot.2013.143.

38. Huppertz, I., Attig, J., D’Ambrogio, A., Easton, L.E., Sibley, C.R., Sugimoto, Y., Tajnik, M., Konig, J., and Ule, J. (2014). iCLIP: protein-RNA interactions at nucleotide resolution. Methods 65, 274–287. 10.1016/j.ymeth.2013.10.011.

39. Van Nostrand, E.L., Pratt, G.A., Shishkin, A.A., Gelboin-Burkhart, C., Fang, M.Y., Sundararaman, B., Blue, S.M., Nguyen, T.B., Surka, C., Elkins, K., et al. (2016). Robust transcriptome-wide discovery of RNA-binding protein binding sites with enhanced CLIP (eCLIP). Nat Methods 13, 508–514. 10.1038/nmeth.3810.

40. Lovci, M.T., Ghanem, D., Marr, H., Arnold, J., Gee, S., Parra, M., Liang, T.Y., Stark, T.J., Gehman, L.T., Hoon, S., et al. (2013). Rbfox proteins regulate alternative mRNA splicing through evolutionarily conserved RNA bridges. Nat Struct Mol Biol 20, 1434–1442. 10.1038/nsmb.2699.

41. Li, W., Chi, H., Salovska, B., Wu, C., Sun, L., Rosenberger, G., and Liu, Y. (2019). Assessing the Relationship Between Mass Window Width and Retention Time Scheduling on Protein Coverage for Data-Independent Acquisition. J Am Soc Mass Spectrom 30, 1396–1405. 10.1007/s13361-019-02243-1.

42. Li, W., Dasgupta, A., Yang, K., Wang, S., Hemandhar-Kumar, N., Chepyala, S.R., Yarbro, J.M., Hu, Z., Salovska, B., Fornasiero, E.F., et al. (2025). Turnover atlas of proteome and phosphoproteome across mouse tissues and brain regions. Cell 188, 2267–2287 e2221. 10.1016/j.cell.2025.02.021.

43. Liu, Y., Mi, Y., Mueller, T., Kreibich, S., Williams, E.G., Van Drogen, A., Borel, C., Frank, M., Germain, P.L., Bludau, I., et al. (2019). Multi-omic measurements of heterogeneity in HeLa cells across laboratories. Nat Biotechnol 37, 314–322. 10.1038/s41587-019-0037-y.

44. Bruderer, R., Bernhardt, O.M., Gandhi, T., Miladinovic, S.M., Cheng, L.Y., Messner, S., Ehrenberger, T., Zanotelli, V., Butscheid, Y., Escher, C., et al. (2015). Extending the limits of quantitative proteome profiling with data-independent acquisition and application to acetaminophen-treated three-dimensional liver microtissues. Mol Cell Proteomics 14, 1400–1410. 10.1074/mcp.M114.044305.

45. Bruderer, R., Bernhardt, O.M., Gandhi, T., Xuan, Y., Sondermann, J., Schmidt, M., Gomez-Varela, D., and Reiter, L. (2017). Optimization of Experimental Parameters in Data-Independent Mass Spectrometry Significantly Increases Depth and Reproducibility of Results. Mol Cell Proteomics 16, 2296–2309. 10.1074/mcp.RA117.000314.

46. Salovska, B., Zhu, H., Gandhi, T., Frank, M., Li, W., Rosenberger, G., Wu, C., Germain, P.L., Zhou, H., Hodny, Z., et al. (2020). Isoform-resolved correlation analysis between mRNA abundance regulation and protein level degradation. Mol Syst Biol 16, e9170. 10.15252/msb.20199170.

